# Subspace alignment as a mechanism for binding

**DOI:** 10.1101/2021.07.07.451472

**Authors:** Justin M. Fine, Seng Bum Michael Yoo, R. Becket Ebitz, Benjamin Y. Hayden

## Abstract

To choose between options, we must solve two important binding problems. First, the features that determine each options’ values must be appropriately combined and kept separate from the corresponding features of other options. Second, options must be associated with the specific actions needed to select them. We hypothesized that the brain solves these problems through use of aligned (for bound dimensions) and orthogonal (for separated dimensions) population subspaces. We examined responses of single neurons in six putative value-coding regions in rhesus macaques performing a risky choice task. In all areas, single neurons encode the features that define the value of each option (stakes and probability) but only very weakly encode value *per se*. However, the coding dimensions associated with these features are aligned on a single subspace, from which a strong emergent value signal can be read out. Moreover, all six regions use nearly orthogonal subspaces for the left and right options, thereby linking options to their position in space, implementing functional partitioning, and reducing the possibility of misbinding. These results provide a new solution to the neuroeconomic binding problems and suggest that other forms of binding may work through similar principles.

## INTRODUCTION

The *binding problem* is one of the classic problems in cognitive neuroscience (Treisman and Gelade, 1980; Treisman, 1996; von der Malsburg, 1995; Roskies, 1999). It deals with the question of how different elements of objects are bound neurally to form a coherent whole. Although not always called that, the binding problem is a central puzzle in decision-making and neuroeconomics as well (Hayden and Moreno-Bote, 2018). Choice options are often defined by multiple independent features, such as probability and stakes in risky choice. Humans and other animals can readily bind these features together to create a single value dimension so that we can compare different combinations of features or bundles of goods (Raghuraman and Padoa-Schioppa, 2014; Pastor-Bernier et al., 2019; Kable and Glimcher, 2007; Kim et al., 2008; Hunt et al., 2014; Kennerley et al., 2009; O’Neill and Schultz, 2010; Azab and Hayden, 2020). Economic choice also often involves a second binding problem, linking offer to action (Wunderlich et al., 2009; Hare et al., 2011; Cai and Padoa-Schioppa, 2014; Hunt et al., 2018). Specifically, we must link the outcome of the value comparison process, which is necessarily abstract, to the specific actions needed to select the option, even if that action is as simple as a reach or a saccade (Rangel et al., 2008; Kable and Glimcher, 2009). Thus, when choosing, brains need to solve feature and action binding problems; failure to successfully solve these problems will inevitably lead to errors in evaluation and choice.

How might the brain solve these binding problems? For perceptual binding, previously proposed solutions include binding by temporal synchrony and binding by specialized neurons that encode multiple variables (Zeki, 2020; Di Lollo, 2012; Von Der Malsburg, 1995 and 1999; Singer and Gray, 1995; Dong et al., 2008). Both solutions have attracted a good deal of criticism, and the problem is still generally considered to be unsolved (e.g., Shadlen and Movson, 1999; Ghose and Maunsell, 1999; Wolfe, 2012; Holcombe and Clifford, 2012). For the neuroeconomic binding problems, solutions have generally focused on specialized abstract value-specific and value-action neurons (e.g., Platt and Glimcher, 1999; Rustichini and Padoa-Schioppa, 2015; Samejima et al., 2005; Cai and Padoa-Schioppa, 2014).

A great deal of single unit physiological research in neuroeconomics is predicated, either explicitly or implicitly, on the idea that neurons are specially tuned for single variables. However, much recent evidence supports the idea that neurons throughout the prefrontal cortex (and likely beyond) exhibit mixed selectivity, or simultaneous, often non-linear encoding for multiple variables (Rigotti et al., 2013; Fusi et al., 2016; Raposo et al., 2014; Blanchard et al., 2018). Mixed selectivity both offers a challenge for traditional interpretational strategies of physiological data and offers an opportunity for new solutions to classic problems in neural coding, such as the binding problem(s). In particular, mixed selectivity can bestow on populations the ability to use high dimensional geometry, which creates a ‘blessing of dimensionality’ and gives room to create specific subspaces for options, into which their component features can be combined, and which can be easily partitioned from other options (Fusi et al., 2016; Cohen et al., 2021; Bernardi et al., 2020; Tang et al., 2020; Parthasarathy et al., 2019; reviewed in Ebitz and Hayden, 2021 and Urai et al., 2021). Moreover, such combinatorial tricks are highly flexible, readily changeable on a moment to moment basic, and easily decoded.

Here, we build on the implications of mixed-selectivity for coding decision-relevant variables and propose a novel population-coding solution to the neuroeconomic binding problems. We propose that the brain may take advantage of population subspaces to combine and partition information. The term subspace refers to a constrained region of the larger space of possible conjoint firing rate patterns for a population of neurons (Gallego et al., 2017; Gao and Ganguli, 2015; Ebitz and Hayden, 2021). Specifically, we propose that populations can carry offer-value information for decisions by aligning subspaces for the features that contribute to offer value. In addition, we propose that the separation of distinct offers can be solved by neural populations making use of separate subspaces. Moreover, these subspaces can be aligned to action-specific dimensions, namely those associated with the action of selecting the option, thereby implementing option-action binding. In summary, subspace alignment offers a ready solution to multiple binding problems.

We recorded responses of neurons in six brain areas associated with reward, the central and rostral orbitofrontal cortices (OFC 13 and rOFC 11), the pregenual and posterior cingulate cortices (pgACC 32 and PCC 29/31), ventromedial prefrontal cortex (vmPFC 14) and the ventral striatum (VS). All regions were recorded in an identical risky choice task involving two options separated in space (left and right) and both defined by probability and stakes, which were selected randomly and were thus uncorrelated (Strait et al., 2014). We found that neurons in all regions encode both variables but use nearly orthogonal subspaces for options appearing on the left- and right-hand side of the screen. We also found, using more conservative analyses than past studies have, that explicit encoding of the expected value (EV) of offers is low at the single neuron level in all regions. But within each spatially partitioned subspace, probability and stakes exist within a single integrative subspace. These results imply that expected value is represented most prominently in an implicit manner through population alignment of feature subspaces. Together, these results endorse the idea that the brain makes use of subspace orthogonalization to partition variables that need to be separated and use subspace alignment to keep together variables that need to be bound.

## RESULTS

### Subjects show unremarkable behavior consistent with task understanding

We examined data collected from six rhesus macaques trained to perform a two-option gambling task that we have used several times in the past (e.g., Strait et al., 2014, **Figure 1**). Data were consistent with patterns we have previously observed (for the most complete analyses, see Farashahi et al., 2018 and 2019). Briefly, all data came from an **asynchronous gambling task**. In this task, macaque subjects use saccades to select between two risky options that appear on the left- or right-hand side of the computer monitor and are presented with a one second asynchrony. Options were defined by a specific probability (0-100%, 1% increments) and stakes (large or medium reward, 0.240 and 0.165 mL juice). The probabilities and stakes for offers 1 and 2 were chosen randomly for each target. The order of presentation was randomized by trial. All subjects showed behavior consistent with task understanding and have been published in past studies, so they are not repeated here. Specifically, preferences depended positive on both probability and stakes, and subjects chose the higher EV option on most trials (Strait et al., 2014 and 2015; Wang et al., 2020; Maisson et al., 2021).

**Figure 1.**
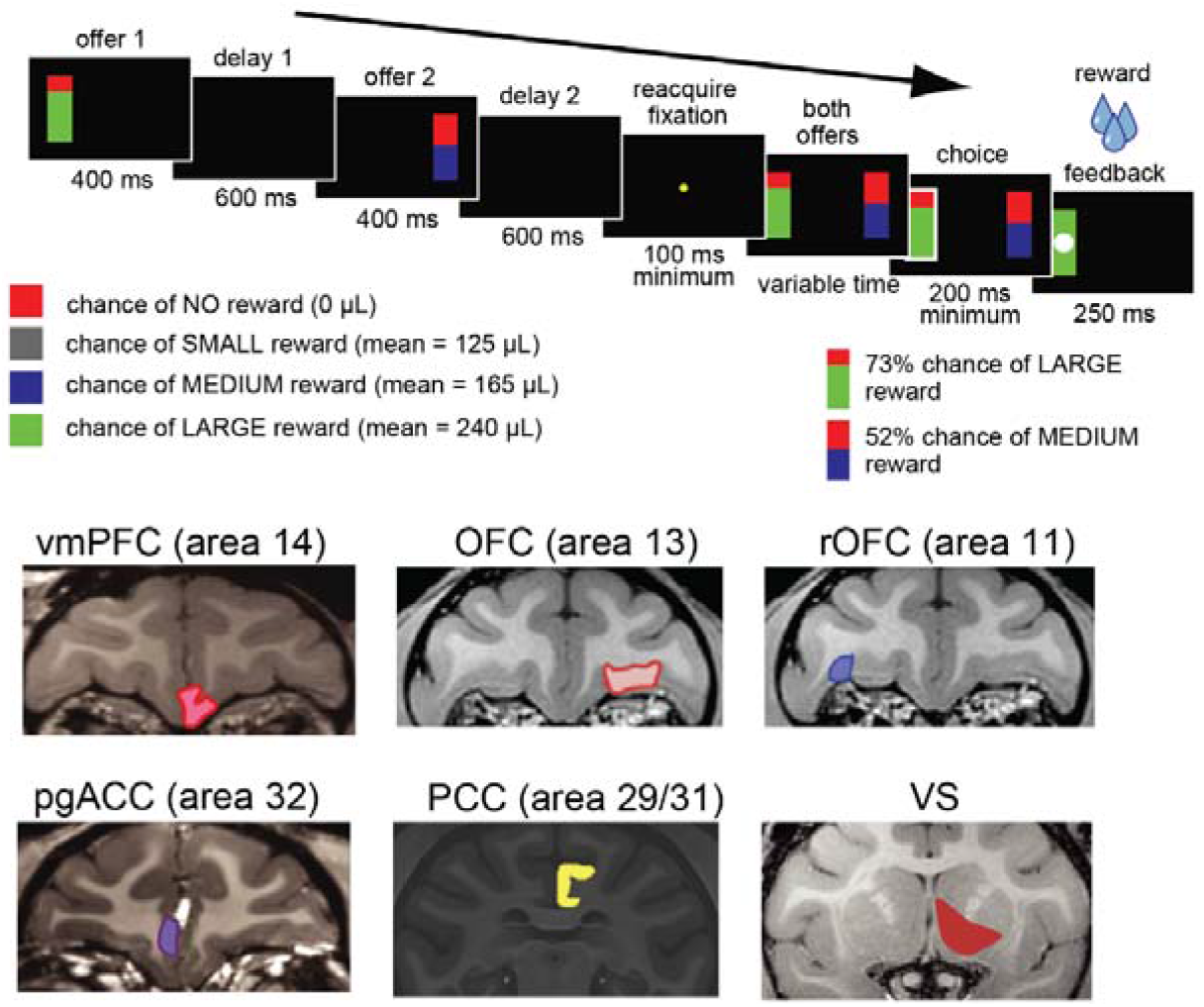
Task outline and brain areas. **a.** The risky-choice task is a sequential offer decision-task that we have used in several previous studies (Strait et al., 2014). In the first 400 ms, subjects see the first offer as a bar presented on either the left or right side. The above shows an example on the left. This offer is followed by a 600 ms delay, a 400 ms offer 2 window, another delay (2) window, and then choice. The full task involved either small, medium or large reward offers. The small reward trials were actually those with safe (guaranteed) offers. We only analyzed the risky choice trials, which were those including the medium and large reward. **b.** MRI coronal slices showing the 6 different core reward regions that were analyzed.

### Pure value coding is rare in single neurons in all areas

At any given moment, a particular neuron may encode any number of decision-relevant features and their interactions - i.e., it may have linear and nonlinear mixed selectivity. To determine how neurons in core value regions code relevant variables, we examined responses of single neurons in six reward coding brain areas (**Figure 1**, OFC 13, rOFC 11, vmPFC 14, pgACC 32, PCC 29/31, and VS, see **Methods**). We recorded a total of 982 neurons (157 in OFC, 138 in rOFC, 156 in vmPFC, 255 in pgACC, 152 in PCC, and 124 in VS). We used a conservative approach that takes into account some of the statistical issues that arise in a world of mixed selectivity. Specifically, unlike most past approaches, our analysis considers the risk of conflating mixed selectivity with pure value coding. We used an elastic-net generalized linear modeling (GLM) approach (see **Methods**); these GLMs included features (probability and stakes) and their interaction. Elastic-net approaches allow us to perform variable selection and estimation simultaneously, while handling potential collinearity between variables (Zou and Hastie, 2003). We quantified the presence of pure value coding at the level of the single neuron. Using the GLM outputs, we counted the number of neurons for which the value coefficient (the interaction of probability and stakes) was non-zero. We performed this analysis separately for both offers.

The proportion of pure expected value neurons did not differ between offer windows either within or across areas; therefore, we present here the numbers as averaged across the two offers and offer epochs. In vmPFC, for example, the average proportion of neurons showing pure coding for value was low, approximately 11.2%. This proportion was the highest of all the regions; the rest of the brain areas had, respectively: OFC, 6.3%; rOFC: 6.8%; pgACC: 7.4%; PCC: 8.5%; VS: 10.2%. Not only was pure value coding rare, but the majority of neurons that did show pure value coding for the value of the first offer were not the ones that showed encoding for the value of the second offer. In other words, it appears that the pure value coding we did observe did not result, for the most part, from specialized pure value neurons. Averaging across brain areas, only 30% (sd 10.4 %) maintained an EV coding between offers, which, given the low numbers of value coding, turns out to be 1-2 neurons per area.

The proportions of “pure” value neurons we find are much lower than those measured in other studies, including our own (e.g., Strait et al., 2014 and 2015). However, our elastic-net approach is a novel approach that avoids the problem of using model-comparison metrics typically used to classify EV neurons. The model-comparison approach can artificially inflate estimates of pure value coding due to collinearity, with the fewer number of variables. In other words, when predictors are correlated, the model selection process will favor a model with singular expected value terms and discard legitimate contributions from the features.

In contrast to the low levels of pure value coding, the proportions of pure value neurons are substantially (and significantly) lower than the proportions showing mixed selectivity (either a linear or nonlinear). Using a test to compare the proportion of pure value and mixed selectivity neurons within each monkey and brain area, we confirmed the proportions of mixed selectivity neurons were higher than pure value (mean *χ^2^(1) = 20.32*, all p < 0.001). Those proportions of mixed selectivity neurons were: vmPFC: 29.1%; OFC: 25.1%; rOFC: 29.1%; pgACC: 24.7%; PCC: 23.1%; VS: 37.8%. These results were consistent over different settings of elastic-net regularization, which we considered the cost mixing parameter ranging from 0.1 (close to Ridge) to 0.9 (close to Lasso), indicating that our results are not an artifact of the specific way we performed our GLM (other data are similar and are therefore not shown). Overall, these results indicate that while pure value coding is rare, mixed selectivity, which most previous studies do not distinguish from pure value coding, is relatively common. That in turn raises the possibility that pure value coding is found predominantly at the population, rather than the single neuron, level.

### Reward regions bind value-relevant features through aligned subspaces

The relative paucity of neurons explicitly encoding value does not, *prima facie*, mean that the brain lacks a way of combining features that constitute value. Indeed, the dependence of behavior on both factors simultaneously implies that such information must be available to the motor system. The alternative we consider here is that the population can bind and integrate features through alignment of feature subspaces, thereby encoding value implicitly (Elsayed et al., 2016; Mante et al., 2013). To test our hypothesis, we examined the geometry between coding dimensions for the two features. We first found a low-dimensional, static population encoding subspace over the first and second offer encoding epochs for each of the features. To do this, we computed the individual feature-encoding subspaces using a standard GLM (rather than elastic-net, see **Methods** and **Figure 2**) that included probability and stakes, EV, and their interactions with spatial location of offer presentation (i.e., left or right). The main reason for using standard GLM here is that standard GLM provides unbiased coefficient estimates, and our focus here was the population distribution rather than the single unit encoding *per se*. We subsequently confirmed that use of elastic-net produces similar results, and, indeed, does not affect any of our statistical outcomes or conclusions (data not shown). The low-dimensional subspaces for the probability and stakes were then found by entering the GLM coefficients (that is, betas) into a singular-value decomposition (SVD) to separate temporal basis and neuron weights (right-hand singular vectors). Note that the SVD was performed separately for probability and stakes, offer 1 and offer 2, and was done over the whole offer epoch 500 ms time windows. We then computed the integrative EV subspace composed of features by correlating the neuron weights from the first SVD dimension of each feature (see **Methods** and **Figure 2**).

**Figure 2.**
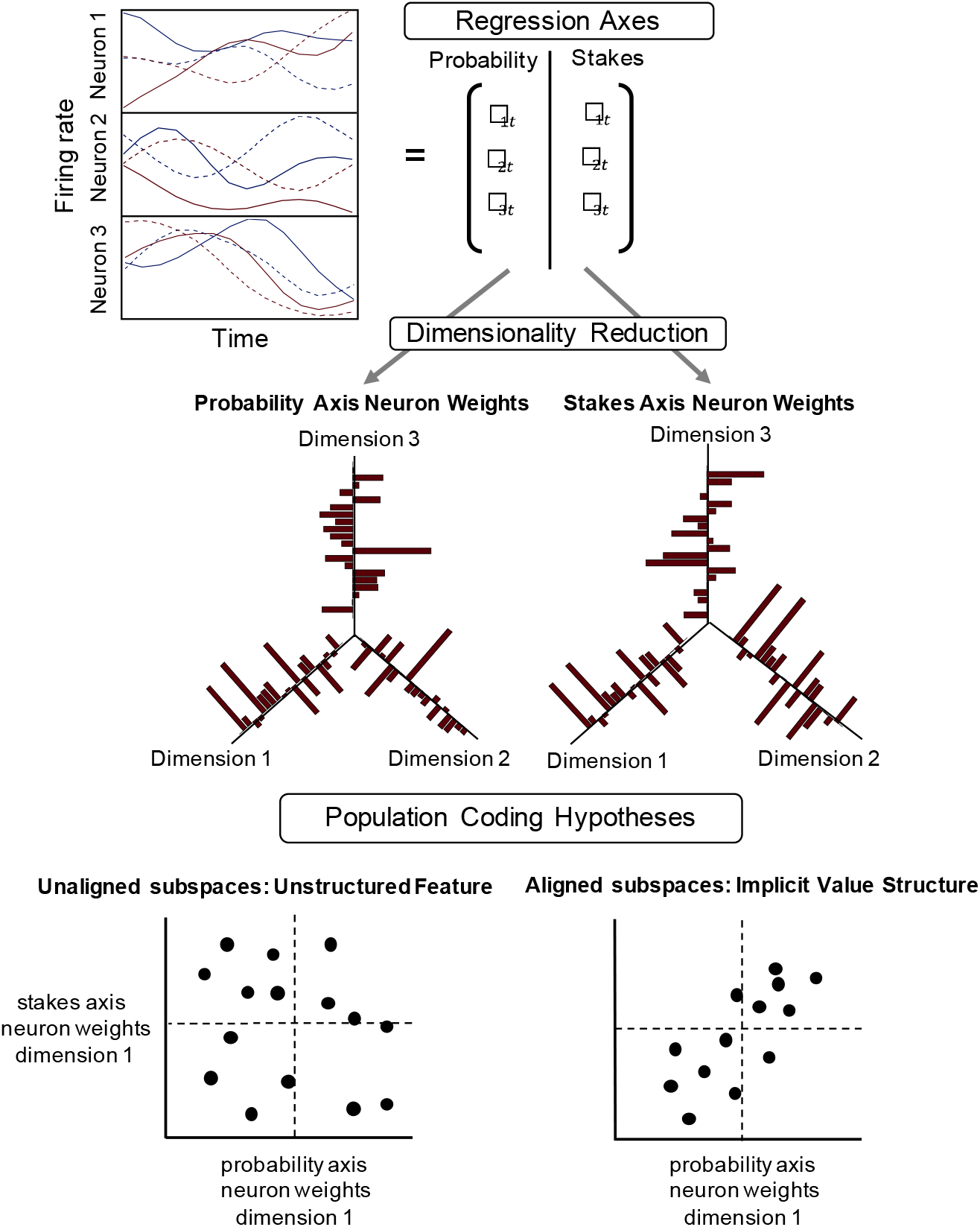
Process of Regression and Dimensionality reduction for obtaining encoding subspaces. to measure feature subspace alignment. **Top.** Panel left shows averaging firing rate of 3 different neurons from OFC (smoothed 90 ms Gaussian). across 3 different levels of probability (colors). The encoding subspaces are obtained using a GLM to obtain the beta parameters for each neuron and time-point. **Middle.** Each variable of probability and stakes is separately entered into a singular value decomposition to obtain neuron weights across the top 3 dimensions. The SVD provides the neural weights for the dominant mode of activity over time in each variable, while separating the temporal bases (not shown). **Bottom.** Different population coding hypotheses. Each of these scatters plots the neuron weights from the 1st dimensions of the preceding SVDs for probability against stakes. The null hypothesis on the left, ‘Unaligned subspaces’ predicts that if value coding is most prominent at the single-unit level, the multivariate distribution of probability and stakes weights will be uncorrelated. Our main hypothesis (displayed on the right), ‘Aligned subspaces’, predicts that population can embed an implicit computation of value by positively aligning the subspaces for probability and stakes. This effectively moves the computation of value away from being explicit in the single neuron. Rather, the population represents it as a scaling of probability responses that depend on reward across the distribution of population basis scaling and mixed-selectivity tuning.

The main hypothesis we consider here is that populations can implicitly code value, which makes a specific prediction regarding how the encoding subspaces for probability and stakes should be distributed. Namely, if the population allows readout of an implicit value code by a downstream observer, it should form a feature integration subspace. This predicts the encoding subspaces for probability and stakes should be non-orthogonal and positively correlated (**Figure 2**, ‘Population Coding Hypotheses’ Right). Otherwise, the ensemble could be organized around a non-integrative and separation of feature codes; in effect, the encoding of probability and stakes will not be correlated (Figure 2, ‘Population Coding Hypotheses’ Left). To control for the possibility that observed correlations are artificially inflated by common trial-specific or noise fluctuations we employed a bootstrapping procedure over the correlations (see **Methods**).

We find that the correlations between probability and stakes subspaces are all non-zero (that is, that they are non-orthogonal). Moreover, they did not differ in magnitude between left or right offer presentation. Indeed, scatter plots of spatial and feature betas for each subject and brain area show that there is a clear positive correlation across areas, which range from 0.55 to 0.72 (**Figure 3A**). Critically for our arguments, all these correlations were well above zero (all *p* < 0.001). This result indicates that, in the population, subspaces for probability and stakes - the two determinants of value - are semi-orthogonal. That is, they are neither perfectly aligned nor fully orthogonal. Our observation of semi-orthogonality suggests the two axes, one for stakes and one for probability, are partially aligned to create a single subspace where the axes are integrated for an implicit computation of the offer-value. The greater alignment implies more integration between the information conveyed by the two feature axes. If they were perfectly collinear, a downstream observer would not be able to read out separate feature contribution. In contrast, the imperfect alignment implies that the individual feature properties are still partially distinguishable downstream.

**Figure 3.**
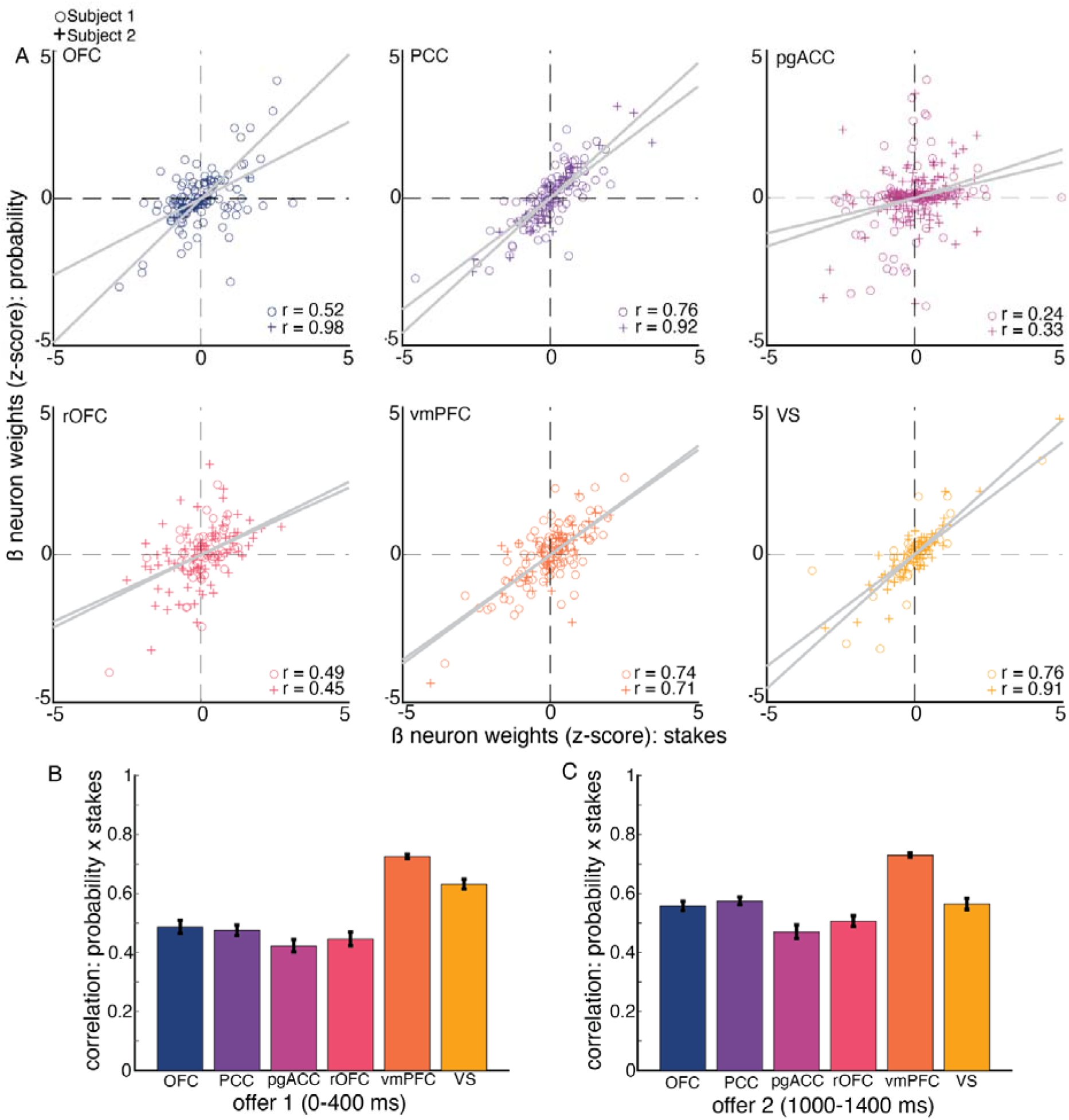
Implicit population codes for value emerge from feature subspace alignment **A.** The scatter plots show the population distribution of probability and stakes neuron weights for each brain and each subject (circles and plus symbols), and linear-squares line of best fit for offer 1. **B** and **C.** Mean bootstrapped correlation coefficients between neuron weights for probability and stakes, and their 95% confidence intervals. Each bar represents a different brain area, which are averaged over both subjects within a brain area. Bars on the left (**A**) and right (**B**) are the correlations for offer 1 and 2 windows, respectively.

### Binding options to specific locations with orthogonal subspaces

How are feature spaces partitioned in space to ensure that options can be distinguished? A viable mechanism would be for populations to use separate subspaces (**Figure 4A**). Subspace separation could be implemented by a conjunctive code that makes use of nonlinear mixed selectivity (Fusi et al., 2016; Bernardi et al., 2020; Tang et al., 2020). Conjunctions of the feature integration subspace and offer location could effectively partition the low-dimensional integration subspace into separable subspaces. We tested this hypothesis by computing a low-dimensional feature integration subspace spanned by both probability and stakes. To do this, we used a tensor-based singular value decomposition (see **Methods** and Tucker, 1966). This method gives the common low-dimensional subspace for the previously computed GLM coefficients for both probability and stakes simultaneously, along with the corresponding neuron weights and temporal basis; essentially, it results in the set of neuron weights for the integrative subspace rather than the features separately. In brief, the tensor or higher-order SVD generalizes the standard SVD from n=2 matrix of variables to an n-dimensional tensor. The tensor SVD returns linear basis vectors that are composed by the sum of its parts, just as in a standard SVD.

**Figure 4.**
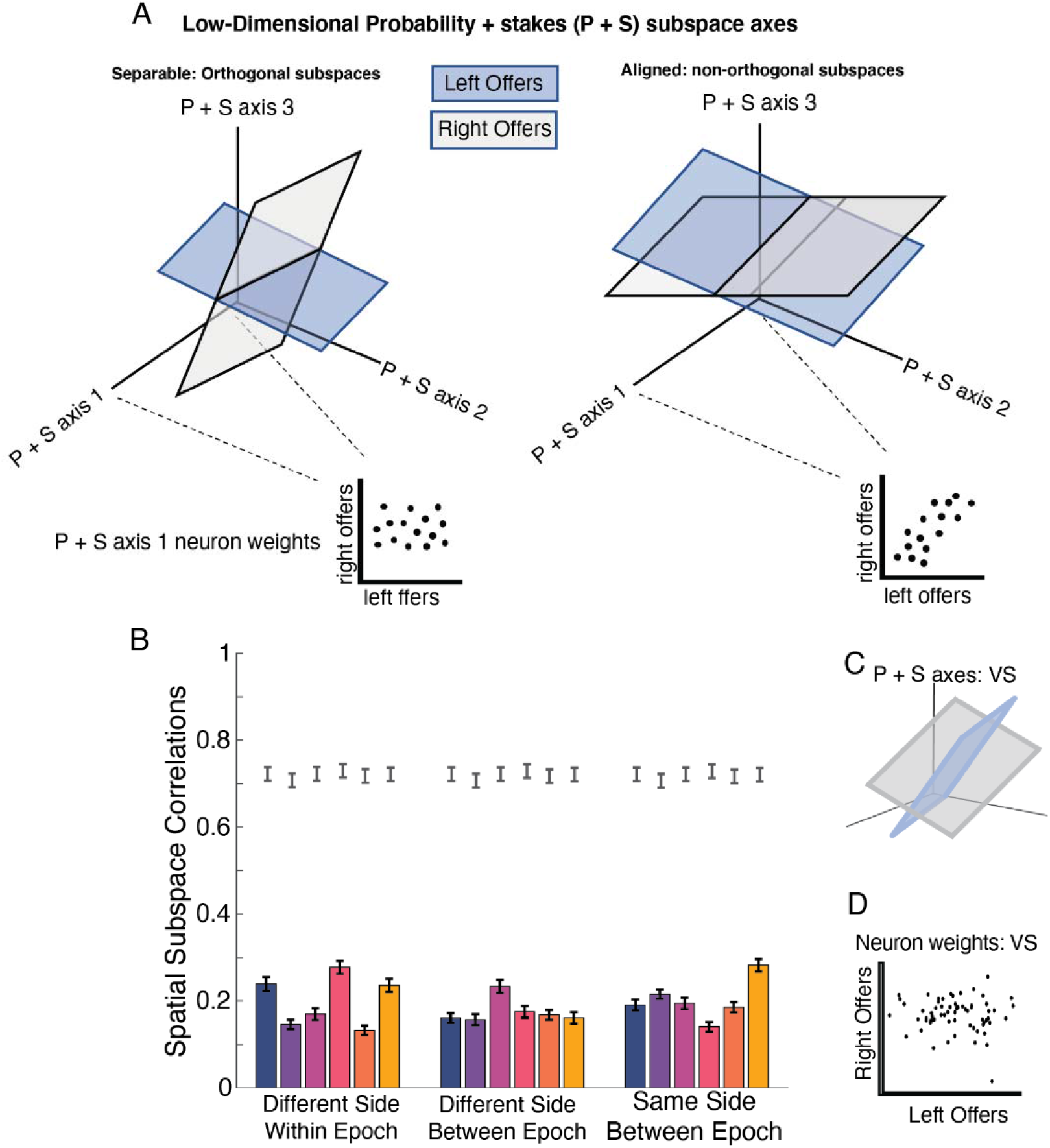
Subspace separability across space can solve binding in neural populations. **A.** The left and right plot show two examples of how an integrative subspace linearly combining probability and stakes (P+S) could be demarcated based on spatial location. In each of these plots are cartoons that depict different outcomes. On the left is our main hypothesis that non-linear mixed selectivity can lead to separable or orthogonal subspaces for different spatial locations of offer presentations. In this scenario, the two planes are approximately 90 degrees apart in 3D space. Our hypothesis can be tested by comparing the neuron weights from dimension 1, obtained from the tensor SVD applied to obtain the P+S low-dimensional space. The cartoon scatter on the left indicates we hypothesize essentially zero correlation between P+S axis 1 weights between offers on the left versus right. As a null hypothesis (**A.** right side), the left and right subspaces may be closely aligned and not present an identifiable mechanism for offer-value binding to space. **B.** Bar plot showing the bootstrapped observed correlations between different neuron weights of different spatial subspaces of offers and their 95% bootstrapped confidence interval. Note, the main hypothesis was that these would all be less than 1 plus noise; the graph shows indeed all were less than 1. The region of significance is denoted by all bars falling outside the 95% Null. The null confidence intervals are displayed as error-bars above each bar. In **B. Different Side: Within**, this is average of offer 1 and 2, comparing left and right within each offer. **B. Different Side: Between**, is the average of comparing left and right between offer 1 and offer 2; for example, one comparison is offer 1 left and offer 2 right. **B. Same Side: Between**, this is the average of comparing the same side, but across offer 1 and 2 epochs; for example, the bars include comparisons of left offers for offer 1 and 2. **C.** An example from a VS subject for the same side between epoch comparison, showing the mean observed 3D subspace plot showing the planes are orthogonal for left and right presentations. **D.** Neuron weights from the same VS subject and condition as **C.** for left and right offers from the first dimension of the P+S axis.

We applied the tensor SVD to find the neuron weights for the integrated probability and stakes subspace. To do this, we used a data tensor of n=3 that included neurons, time, and a variable mode that includes both probability and stakes GLM coefficients. The variable mode will yield a weight for each of probability and stakes, which we refer to as the probability + stakes mode (which we will call P + S). Therefore, we computed the n=3 tensor SVD separately for left and right offers to get their subspace neuron weights. To determine separability, we correlated the neuron weights from the first dimension of each tensor SVD for different spatial locations pairwise.

To create a criterion for what is considered separable, we first generated a null sampling distribution or “upper ceiling” perfect correlation that would occur when only corrupted by noise (see **Methods**). The null correlations were computed by bootstrapping the probability and stakes subspaces and correlating them with themselves. We compared the 99% confidence intervals from the null distribution against the observed correlations to determine separability. In addition, we bootstrapped the observed correlations to determine confidence intervals and in doing so, computed the distance from the null. We also compared our observed correlations to chance (that is, to zero, p<0.05).

When comparing subspaces between different sides (left and right), the null correlations (i.e., the observable ceiling) for each brain area had absolute values ranging from 0.74 to 0.88 (lines in **Figure 4B**). All of the observed estimates of subspace separability were well below these values (bars in **Figure 4B**). Specifically, all brain areas and all comparisons exhibited a large degree of separability from the theoretical lower bound established by the null confidence intervals (all *p* < 0.001), and they were also greater than zero (*p* < 0.001, **Figure 4B**). Additionally, there were no notable differences within an area or across comparisons. By demonstrating here that the feature interaction subspaces are separable or orthogonal with respect to space indicates that the brain makes use of spatial position to organize distinct regions of the entire manifold, allowing it to maintain different offers with minimal overlap.

### Like offers, chosen options are integrated and segregated by means of subspaces

After offers are presented, decision makers can choose a preferred one. Some findings support a distinction between how offered and chosen values are encoded (e.g., in separate specialized populations of neurons, as in Padoa-Schioppa and Assad, 2006). We therefore next asked whether the population uses similar or different principles for binding and separation for chosen offers. For these analyses, we focused on the 400 ms time-window immediately preceding the expression of choice (by means of saccade), a time during which the screen was blank. Our focus on the blank period is motivated by a desire to reduce stimulus-driven effects from neuronal responses. One possibility is that chosen options obey the same coding principles as offered ones; another is that latent or implicit value coding becomes explicit on being chosen.

We find that population encoding for chosen option features follow the same principles as encoding for offer features. As in our analysis of offer values, we used a GLM with elastic-net regularization to determine the proportion of the population exhibiting significant encoding of chosen value. Much like the coding of offer value, there was a weak showing of chosen value at the single neuron level across all areas; OFC showed 11.2% encoding; the other regions had a similar, and indeed, smaller proportion: vmPFC: 6.3%; rOFC: 3.2%; pgACC: 3.8%; PCC: 2.9%; VS: 3.1%. As above, these results were consistent over different settings of regularization (data not shown). These proportions are substantially lower than the proportions showing mixed selectivity (either a linear- or nonlinear-mixed selectivity) across all regions (vmPFC: 32.5%; OFC: 25.4%; rOFC: 26.1%; pgACC: 27.1%; PCC: 30.1%; VS: 19.8%), which were also confirmed using a chi-square test (mean *χ^2^(1) = 18.42*, all p < 0.001). These results indicate that mixed selectivity is the dominant response mode of chosen value coding.

As with offered value, we found strong latent encoding of chosen value in the ensemble response. Specifically, we repeated our approach from above, this time using chosen rather than offered value, and again, bootstrapped the GLM coefficients to minimize trial-specific fluctuations. The GLM included chosen probability and chosen stakes, chosen EV, chosen side of offer and all interactions. We used the same SVD approach as above to find separate neuron weights for probability and stakes and correlated neuron weights to determine the extent of feature integration for chosen options. We found that correlations between chosen probability and chosen stakes subspaces were all non-zero (all *p*<0.001) and did not differ in magnitude between left or right offer choice. Specifically, they ranged from 0.23 to 0.74 across brain areas, albeit with modest inter-region variation in correlation: vmPFC: 0.57; OFC: 0.35; rOFC: 0.60; pgACC: 0.28; PCC: 0.46; VS: 0.48. In other words, all six regions used aligned subspaces to implement binding across dimensions of shared features of common objects.

As with offers, chosen options are linked to specific positions in space through subspace separation. We used the same technical approach here that was employed above for comparing spatial separation in offer value subspaces. Namely, we projected chosen probability and stakes coefficients to common subspace through tensor SVD. We computed separate decompositions for left and right chosen offers, controlling for chosen offer order (1st or 2nd) in the GLM, and then proceeded to correlate the neuron weights. Again, the observed correlations were calculated through bootstrapping and compared to a null distribution. All the subspace correlations comparing space were below the 95% confidence intervals established by the null distributions. Across regions the observed correlations ranged from 0.12 in OFC to 0.43 at the largest in VS. The rest of the regions showed intermediate values: vmPFC: 0.17; rOFC: 0.36; pgACC: 0.14; PCC: 0.14. These results indicate that, as with offered options, features of chosen offers occupy separate subspaces according to the spatial location of the chosen offer. In other words, all six reward regions use different codes for the features of chosen left offers and chosen right offers.

## DISCUSSION

When we perceive an object, we must bind its distinct features to form a subjective whole (Von Der Malsburg, 1981 and 1999; Roskies, 1999; Treisman, 1996). Understanding this process, that is, solving the ***binding problem***, is important in perception. The question of how features are integrated to a whole is also important, although not always under that name, in neuroeconomics (Hayden and Moreno-Bote, 2018). Here we propose a new solution to the neuroeconomic binding problems, one that also holds promise for perceptual binding. Specifically, we propose that binding and partitioning may occur via subspace alignment and separation. We then provide novel unanticipated physiological results consistent with that solution. Our results support the idea that the brain makes use of separate coding axes for distinct feature dimensions, and, to integrate features, combines them into a single subspace that spans the two coding axes. The objects represented in that subspace must then be separated or partitioned from other objects to ensure effective choice and to avoid misbinding features of different objects. We propose that this partitioning problem can be solved by use of different subspaces for objects; one natural organizing principle would be to bind these dimensions to specific positions in space, such as those in which the objects are located. Indeed, that particular partitioning scheme serves a second purpose, namely, the binding of the object to the information about the action needed to select it, and thereby solves a separate action binding problem.

Our proposal makes central use of subspaces, which are afforded by ensemble coding (Fusi et al., 2016; Bernardi et al., 2020; Ebitz and Hayden, 2021; Urai et al., 2021). Ensemble coding allows for multiple neurons to contribute to a single representation, thus affording a ‘blessing of dimensionality’. Indeed, each additional neuron that contributes a new nonlinear mixture or conjunction code to the population can greatly expand the expressiveness of the coding space (Fusi et al., 2016). That expansion, in turn, creates room to separate different objects in different subspaces; each subspace can be shared by features of the same object. More generally, subspace alignment and orthogonalization, our proposed solution to the binding problems, is a powerful tool for segregating information. For example, it is used to separate motor planning and motor execution (Elsayed et al., 2016) and economic evaluation from selection (Yoo and Hayden, 2020). One convenient feature of using subspaces for partitioning is that they are highly flexible and can rapidly adapt to changing circumstances; they therefore allow for rapid flexible choice under changing circumstances. Indeed, a felicitous feature of our proposal is that subspace partitioning can fail (see, for example, Tang et al., 2020). Far from being a limitation, the ability to explain binding failures is an essential feature of any successful theory of binding (Wolfe, 2012; Holcombe and Clifford, 2012).

Perhaps the most surprising finding we present is that the location in which an offer occurs affects not just the firing of the neuron that encodes its features, but also the way in which the population of neurons encode those features. In other words, space alters firing, but does not simply increase or decrease it; instead, space alters its geometry. Our results therefore demonstrate the limitations of previous proposals, including our own, which imagined space as a modulatory factor that regulated the gain of value coding (Strait et al., 2016; Yoo et al., 2018; Roesch and Schoenbaum, 2006; Feierstein et al., 2006; Tsujimoto et al., 2009). Historically, lack of spatial tuning has been taken to be a hallmark of abstract or pure value coding. However, the repeated demonstration of spatial effects on responses of reward neurons has challenged this idea. Our results here advance the story further and indicate that space can be used as an organizing principle for specialized subspaces that contributed to binding and partitioning.

We propose that, in our dataset, spatial location serves as the binding anchoring point for subspaces. However, decision-making over options is not specific to space. One advantage of subspace partitioning is that it does not need to rely on space as an organizing principle; it can extend to more seemingly abstract dimensions. For example, economic choice often involves deliberation between options in which space is unspecified (for example, a waiter naming three desserts and asking for a preferred one). We hypothesize that in such cases, partitioning will occur through alignment to some other non-spatial dimension. For example, if options are separated in time, the organization may be time-based (indeed, we show this is possible, see **Results**). More generally, we surmise that the brain may make use of any organizing scheme at hand to define subspaces and then use those to implement partitioning and avoid misbinding.

Our GLM analysis shows a relatively small number of neurons with pure value coding in any of these regions; instead, we find that mixed selectivity is the norm. This result may appear surprising given the large literature showing value selectivity robustly in core reward regions, including our own past studies (e.g., Strait et al., 2014). However, our analysis is, as far as we know, the first to strictly follow the surprisingly deep implications of mixed selectivity for theories of neural coding as it pertains to value. Indeed, the present study is the first we know of to exclude demonstrably mixed selective neurons from the possible “pure” value coding. In contrast, work demonstrating greater numbers of pure value neurons tends to be done under the assumption of single selectivity and engages in model selection procedures that wind up conflating mixed selectivity with value coding. That approach will necessarily inflate estimates of pure value encoding in the case of a sample of neurons that shows mixed selectivity. Our results do not, however, indicate that the brain does not encode value, they suggest that value can be readily encoded even if it is not available in single neurons. Indeed, these results accord with recent results from Kimmel et al. (2020), which, using population approaches, show that value and choice are separable from one another. Their results indicate that neurons are not necessarily tuned to chosen value, but likely represent distinct subspaces for choice and value that are integrated in the population. Our results extend these ideas down to the core value signal, indicating that value might also reflect the alignment of compositional feature subspaces. This conclusion is particularly compelling given that a swath of theories that link areas such as OFC with deriving world states or values on-the-fly from features, rather than recalling a stored value (Niv, 2019).

Our results here qualify and, in important ways, disconfirm our previous predictions (Hayden and Moreno-Bote, 2018). In that paper, we argued that if the attentional selection is limited to a single option, then changes in neural activation must be related to that option, its features, and its afforded actions. Our results here indicate that that past prediction was, at best, incomplete because it only accounts for cases in which options are processed serially. By contrast, subspace orthogonalization allows for decision-makers to keep two objects in mind at the same time and minimize interference between their encodings. We do not think, however, that our previous proposal was fully wrong – there are clearly many cases in which options are considered serially, even if there are many others in which options are considered simultaneously (Rich and Wallis, 2016; Krajbich et al., 2010; Kacelnik et al., 2011). Likewise, attentional focus often alternates between objects in series, but can in many cases be split between objects (McMains and Somers, 2004; Cavanagh and Alvarez, 2005; Kawahara and Yamada, 2006). Our theory here, therefore, offers a more general accounting of the way the brain binds and partitions information during choice.

These findings across subspace comparisons were, perhaps surprisingly, consistent across all six reward regions we examined and in both members of each pair. This finding indicates that the principles of feature and spatial binding are shared across multiple value-coding regions and raise the possibility that they reflect a general principle by which corticostriatal reward systems represent information about offers and choices. Our results provide evidence, therefore, against modular theories of value coding, ones in which different regions play different nameable functions, such as evaluation, comparison, and action selection (Rangel et al., 2008). We believe instead that they cohere most closely with theories in which economic choice reflects the operation of distributed processes operating simultaneously on distributed representations in multiple regions (Hunt et al., 2014; Cisek, 2012; Pezzulo and Cisek, 2016; Hunt and Hayden, 2017; Yoo and Hayden, 2018; Fine and Hayden, 2021)

## METHODS

### Surgical procedures

All procedures were approved by either the University Committee on Animal Resources at the University of Rochester or the IACUC at the University of Minnesota. Animal procedures were also designed and conducted in compliance with the Public Health Service’s Guide for the Care and Use of Animals. All surgery was performed under anesthesia. Male rhesus macaques (*Macaca mulatta*) served as subjects. A small prosthesis was used to maintain stability. Animals were habituated to laboratory conditions and then trained to perform oculomotor tasks for liquid rewards. We placed a Cilux recording chamber (Crist Instruments) over the area of interest. We verified positioning by magnetic resonance imaging with the aid of a Brainsight system (Rogue Research). Animals received appropriate analgesics and antibiotics after all procedures. Throughout both behavioral and physiological recording sessions, we kept the chamber clean with regular antibiotic washes and sealed them with sterile caps.

### Recording sites

We approached our brain regions through standard recording grids (Crist Instruments) guided by a micromanipulator (NAN Instruments). All recording sites were selected based on the boundaries given in the Paxinos atlas (Paxinos et al., 2008). In all cases we sampled evenly across the regions. Neuronal recordings in OFC were collected from subjects P and S; recordings in rOFC were collected from subjects V and P; recordings in vmPFC were collected from subjects B and H; recordings in pgACC were collected from subject B and V; recordings from PCC were collected from subject P and S; and recording in VS were collected from subject B and C.

We defined **rOFC 11** as lying within the coronal planes situated between 34.05 and 42.15 mm rostral to the interaural plane, the horizontal planes situated between 4.5 and 9.5 mm from the brain’s ventral surface, and the sagittal planes between 3 and 14 mm from the medial wall. The coordinates correspond to area 11 in Paxinos et al. (2008).

We defined **OFC 13** as lying within the coronal planes situated between 28.65 and 34.05 mm rostral to the interaural plane, the horizontal planes situated between 3 and 6.5 mm from the brain’s ventral surface, and the sagittal planes between 5 and 14 mm from the medial wall. The coordinates correspond to area 13m in Paxinos et al. (2008). We used the same criteria in a different dataset (Blanchard et al., 2015).

We defined **vmPFC 14** as lying within the coronal planes situated between 29 and 44 mm rostral to the interaural plane, the horizontal planes situated between 0 and 9 mm from the brain’s ventral surface, and the sagittal planes between 0 and 8 mm from the medial wall. These coordinates correspond to area 14m in Paxinos et al. (2008). This dataset was used in Strait et al., 2014 and 2016.

We defined **pgACC 32** as lying within the coronal planes situated between 30.90 and 40.10 mm rostral to the interaural plane, the horizontal planes situated between 7.30 and 15.50 mm from the brain’s dorsal surface, and the sagittal planes between 0 and 4.5 mm from the medial wall (**Figure 1B**). Our recordings were made from central regions within these zones, which correspond to area 32 in Paxinos et al. (2008). Note that we the term 32 is sometimes use more broadly than we use it, and in those studies encompasses the dorsal anterior cingulate cortex; we believe that that region, which is not studied here, should be called area 24 (Heilbronner and Hayden, 2016).

We defined **PCC 29/31** as lying within the coronal planes situated between 2.88 mm caudal and 15.6 mm rostral to the interaural plane, the horizontal planes situated between 16.5 and 22.5 mm from the brain’s dorsal surface, and the sagittal planes between 0 and 6 mm from the medial wall. The coordinates correspond to area 29/31 in Paxinos et al. (2008, Wang et al., 2020).

We defined **VS** as lying within the coronal planes situated between 20.66 and 28.02 mm rostral to the interaural plane, the horizontal planes situated between 0 and 8.01 mm from the ventral surface of the striatum, and the sagittal planes between 0 and 8.69 mm from the medial wall. Note that our recording sites were targeted towards the nucleus accumbens core region of the VS. This datset was used in Strait et al., (2015 and 2016).

We confirmed recording location before each recording session using our Brainsight system with structural magnetic resonance images taken before the experiment. Neuroimaging was performed at the Rochester Center for Brain Imaging on a Siemens 3T MAGNETOM Trio Tim using 0.5 mm voxels or in the Center for Magnetic Resonance Research at UMN. We confirmed recording locations by listening for characteristic sounds of white and gray matter during recording, which in all cases matched the loci indicated by the Brainsight system.

### Electrophysiological techniques and processing

Either single (FHC) or multi-contact electrodes (V-Probe, Plexon) were lowered using a microdrive (NAN Instruments) until waveforms between one and three neuron(s) were isolated. Individual action potentials were isolated on a Plexon system (Plexon, Dallas, TX) or Ripple Neuro (Salt Lake City, UT). Neurons were selected for study solely on the basis of the quality of isolation; we never preselected based on task-related response properties. All collected neurons for which we managed to obtain at least 300 trials were analyzed; no neurons that surpassed our isolation criteria were excluded from analysis.

### Eye-tracking and reward delivery

Eye position was sampled at 1,000 Hz by an infrared eye-monitoring camera system (SR Research). Stimuli were controlled by a computer running Matlab (Mathworks) with Psychtoolbox and Eyelink Toolbox. Visual stimuli were colored rectangles on a computer monitor placed 57 cm from the animal and centered on its eyes. A standard solenoid valve controlled the duration of juice delivery. Solenoid calibration was performed daily.

### Risky choice task

The task made use of vertical rectangles indicating reward amount and probability. We have shown in a variety of contexts that this method provides reliable communication of abstract concepts such as reward, probability, delay, and rule to monkeys (e.g. Azab et al., 2017 and 2018; Sleezer et al., 2016; Blanchard et al., 2014). The task presented two offers on each trial. A rectangle 300 pixels tall and 80 pixels wide represented each offer (11.35° of visual angle tall and 4.08° of visual angle wide). Two parameters defined gamble offers, *stakes* and *probability*. Each gamble rectangle was divided into two portions, one red and the other either gray, blue, or green. The size of the color portions signified the probability of winning a small (125 μl, gray), medium (165 μl, blue), or large reward (240 μl, green), respectively. We used a uniform distribution between 0 and 100% for probabilities. The size of the red portion indicated the probability of no reward. Offer types were selected at random with a 43.75% probability of blue (medium magnitude) gamble, a 43.75% probability of green (high magnitude) gambles, and a 12.5% probability of gray options (safe offers). All safe offers were excluded from the analyses described here, although we confirmed that the results are the same if these trials are included. Previous training history for these subjects included several saccade-based laboratory tasks, including a cognitive control task (Hayden et al., 2010), two stochastic choice tasks (Blanchard et al., 2014; Heilbronner and Hayden, 2016), a foraging task (Blanchard and Hayden, 2015), and a discounting task (Pearson et al., 2010).

On each trial, one offer appeared on the left side of the screen and the other appeared on the right. We randomized the sides of the first and second offer. Both offers appeared for 400 ms and were followed by a 600-ms blank period. After the offers were presented separately, a central fixation spot appeared, and the monkey fixated on it for 100 ms. Next, both offers appeared simultaneously and the animal indicated its choice by shifting gaze to its preferred offer and maintaining fixation on it for 200 ms. Failure to maintain gaze for 200 ms did not lead to the end of the trial but instead returned the monkey to a choice state; thus, monkeys were free to change their mind if they did so within 200 ms (although in our observations, they seldom did so). Following a successful 200-ms fixation, the gamble was resolved and the reward was delivered. We defined trials that took > 7 sec as inattentive trials and we did not include them in the analyses (this removed ~1% of trials). Outcomes that yielded rewards were accompanied by a visual cue: a white circle in the center of the chosen offer. All trials were followed by an 800-ms intertrial interval with a blank screen.

#### Neural processing, data-selection, and statistical analysis

We calculated firing rates in 60-ms bins but we analyzed them in longer (500 ms) epochs. For estimating GLM coefficients (see below), firing rates were all Z-scored for each neuron. In GLM estimates, we only included risky trials. No smoothing was applied before any reported analysis.

### Single-Neuron encoding classification using elastic net general linear model

To estimate the extent to which neurons could be classified as encoding any number of variable combinations, we used an elastic-net regularized general linear model (GLM; using glmnet package in Matlab) approach with cross-validation. Specifically, our aim was to determine the extent to which neurons could be called ‘pure value’ coding versus a coding of a mixture of other variables, including probability or stakes and their interaction. Elastic-net is a form of regularization that includes both L1 and L2-norms of both Lasso and Ridge regression, respectively. Elastic-net GLMs can provide parameter estimates with reduced bias and variance when there is collinearity in the predictors (Zou and Hastie, 2003). Specifically, the Lasso regularization has the effect of shrinking weakly contributing predictors to 0. The ridge regularization has the effect of reducing the size of all predictors. The elastic-net represents a mixture of both these costs in determining GLM parameters. Importantly, this approach afforded a data-principled approach to identifying the neuron coding.

GLM (regression) coefficients were estimated for the average firing rate during the offer 1 encoding window (1-500 ms) and separately for each neuron to fit the firing rates (FR):

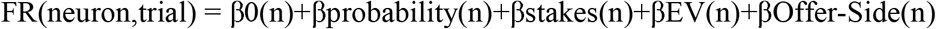

Using the elastic-net GLM has 2 parameters that must be specified for estimation, which include (1) alpha, which specifies the relative impact of L1 (Lasso) and L2 (Ridge) costs, and (2) lamba, the smoothing parameter. Alpha of 0, for example, will give a pure Ridge while alpha of 1 is pure Lasso; alpha in-between these values are the elastic-net or a mixture of costs. Here, we used an alpha of 0.5 to use a balanced elastic-net model that compromised between coefficient retainment and coefficient estimation. To choose the smoothing parameter lambda, we estimated the model over a range of lambdas (100 values log10 spaced from 10° to 10^−3^) using 10 fold cross-validation.

To classify the coding of neurons, the number of neurons coding purely for value versus a mixture was used by counting the non-zero coefficients for each neuron. The lambda with the smallest model deviance was used to select the coefficients for that neuron. Pure value coding was denoted as neurons that only had a singular non-zero βEV coefficient. Neurons exhibiting a mixture of 2 or more variables were all denoted as a mixed-selectivity neuron.

### Dimensionality Reduction for feature subspaces and population EV computation

To determine whether the population ensemble exhibits a scaling of probability and stakes features needed for population level EV encoding, we first computed the individual feature encoding regress subspaces using a standard GLM. We used a standard GLM to estimate the coefficients rather than an elastic-net for two related reasons. First, standard GLM or ordinary least squares – unlike elastic net regularization – is an unbiased estimator and our aim was to recover coefficient estimates for each neuron to compare their distribution at the population level. Specifically, the population analysis was focused on the relation between neuron coefficients across the population whereas the elastic-net was ideal for dealing with collinearity in estimating the effect of predictors in the individual units. Second, and related to the first, using a standard GLM even with collinear predictors at the single-unit level will still only retain unique predictor contributions.

The GLM included predictors for probability and stakes, EV, and their interactions with spatial location of offer presentation (i.e., left or right). The core hypothesis underlying this approach is that, while the individual units may exhibit weak EV coding, the population may do so strongly. Specifically, our hypothesis predicts a positive correlation between GLM coefficients for probability and stakes -- even when accounting for EV presence in the model. The premise of our idea is that the coefficients represent a stable set of feature coding subspace axes that are aligned.

To find the low-dimensional population subspace for each variable, probability and stakes, we entered the β probability(n,t) and β stakes(n,t) coefficients into separate singular-value decompositions (SVD). This was done separately for the offer 1 and 2 time-windows. To determine the extent each area exhibited aligned feature subspaces. Before correlating these neuron weights, we addressed the potential that the alignment of feature subspaces could reflect artificial inflation from trial-specific fluctuations. Therefore, we based our analysis on the assumption that aligned (correlated) neuron weights for both variables should occur regardless of the trial-set used in the GLM.

Effectively, we computed the correlations by bootstrapping the GLM estimation using subsets of trials with replacement. We computed 10,000 GLMS for each neuron and used 70% of randomly selected trials on each bootstrap. The 70% dataset size was chosen heuristically by looking for a size that would exhibit relatively stable coefficient estimates across subsets of trials. This was achieved by performing the same GLM bootstrapping procedure for dataset sizes of 50% to 90% in 5% increments and finding the point where the standard deviation of betas differed between dataset sizes by less than 0.01 on average across coefficients, brain areas and subjects. The motivation for this approach was that the coefficients should have a reliable variability regardless of trials used for estimation. Finally, we ran each bootstrap through the SVD to estimate the neuron weights for each variable, then we computed the correlations between all bootstrap sets. The mean and 95% confidence intervals were calculated using the off-diagonal correlations to avoid inclusion of correlations between those weights derived from the same trials.

### Identifying spatially separable population offer-EV subspaces

To measure the alignment (or orthogonality) between implicit value subspaces across space, we also used a combined GLM and SVD approach, and the correlation of neuron weights. We used the previously computed GLM coefficients from calculating the population EV computation. Here, though, our goal was to compare the implicit EV subspace between offers on the left and right and across offer windows. Our hypothesis was that there are separable EV subspaces for left and right offers, that enable distinct downstream readout of different offers.

To compute the EV subspace as a function of space and offer, we used a higher-order tensor SVD approach. Specifically, we entered the β probability(n,t) and β stakes(n,t) into the SVD together. Entering in both variables together allows to represent the probability and stakes subspace as a linear combination. These weights define a vector in 2D space and are linked to the neuron weights and temporal modes through the outer products of the different modes (see Figure X). What’s important here is that each dimension of a P+S mode (Figure x) has corresponding neuron weights that account for the shared weighting of both variables. In effect, the neuron weights are for an integrated subspace of probability and stakes. The SVD was used separately for left and right for offer 1 and left and right for offer 2. This yielded four different low-dimensional spaces. There are six distinct task-functional comparisons, including comparisons of (1) left and right within offers 1 and 2, (2) left and right between offers 1 and 2, and (3) between the same-side across offers (i.e., left and left for offer 1 and 2, right and right for offer 1 and 2).

The testing of separability between subspaces requires an approach for establishing a null hypothesis of 1. Specifically, because our main hypothesis was essentially a close to zero correlation between subspaces (i.e., |r|=0), we needed to estimate how a perfect correlation in the dataset, but confounded by noise, would be distributed (|r|=1). Therefore, we consider a subspace as separable or (semi-)orthogonal if the correlation between neuron weights is outside the confidence bound of this hypothetical perfect correlation distribution.

We addressed this problem by applying an already established approach (Kimmel et al., 2020), which we describe in brief. They (and Spearman, 1970) show that the effect of noise in correlations can be accounted for in a perfect correlation by estimating the reliability of coefficients and their resultant correlations. Specifically, the null hypothesis of no separability between a left and right space can be written as 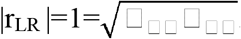. The reliability and accountancy of noise is done using the bootstrapped and tensor SVD derived neuron weights to compute, for example, r_LL_. The element-wise product is then taken between r_LL_ and r_RR_ to compute the sampling distribution under the null hypothesis of perfect correlation influenced by noise. To determine whether the observed correlations r_LR_ was statistically different from the estimated null, we computed the difference between their lower and upper-bound of their 95% confidence interval. Effectively, non-overlap between these two imply at least a *p* < 0.05 statistical difference between the means.

### EV encoding, feature subspace integration and spatial subspace separability for chosen offers

We also analyzed the encoding of activity corresponding specifically to the chosen option. In doing this, we repeated all of the analyses from above but over different time-windows and slightly different predictors (described below). The time-window for analyzing choice was the 400 ms after offer 2 presentation, wherein the screen was blank. As noted in the results, this was chosen to mitigate effects of exogenous stimuli. Namely, it would be impossible to differentiate encoding of offer 2 versus choice of the 2nd offer in a GLM. Analysis during the blank 400 ms window afforded a differentiation since there were no active stimuli.

The other difference in our analysis of choice was the specification of predictors for the elastic-net or standard GLMs. Both the models for choice were generally the same as specified for offer encoding, except that the β’s for probability and stakes, as well as side, referred to the chosen options rather than the specific offers. Additionally, because we were interested in the spatial separation of subspace, we marginalized these results over the distinction between choice of offer 1 or offer 2. Other than these differences, all details were the same.

## Acknowledgements

We thank Bill Vinje, who led an excellent summer journal club on the binding problem at UC Berkeley in 2001. We thank Maya Wang, Tyler Cash-Padgett, Marc Mancarella, Caleb Strait, Tommy Blanchard, and Brianna Sleezer for assistance with data collection.

## Notes

**Funding statement** This research was supported by a National Institute on Drug Abuse Grant R01 DA038615 and MH124687 (to BYH).

### Competing Interest Statement

The authors have declared no competing interest.

### Summary of Updates

Figure 2 was grainy. Revised version is fixed.

## REFERENCES

Azab, H., & Hayden, B. Y. (2020). Partial integration of the components of value in anterior cingulate cortex. Behavioral Neuroscience, 134(4), 296.

Azab, H., & Hayden, B. Y. (2017). Correlates of decisional dynamics in the dorsal anterior cingulate cortex. PLoS biology, 15(11), e2003091.

Azab, H., & Hayden, B. Y. (2018). Correlates of economic decisions in the dorsal and subgenual anterior cingulate cortices. European Journal of Neuroscience, 47(8), 979–993.

Bernardi, S., Benna, M. K., Rigotti, M., Munuera, J., Fusi, S., & Salzman, C. D. (2020). The geometry of abstraction in the hippocampus and prefrontal cortex. Cell, 183(4), 954–967.

Blanchard, T. C., Wolfe, L. S., Vlaev, I., Winston, J. S., & Hayden, B. Y. (2014). Biases in preferences for sequences of outcomes in monkeys. Cognition, 130(3), 289–299.

Blanchard, T. C., Piantadosi, S. T., & Hayden, B. Y. (2018). Robust mixture modeling reveals category-free selectivity in reward region neuronal ensembles. Journal of neurophysiology, 119(4), 1305–1318.

Blanchard, T. C., & Hayden, B. Y. (2015). Monkeys are more patient in a foraging task than in a standard intertemporal choice task. PloS one, 10(2), e0117057.

Blanchard, T. C., Wilke, A., & Hayden, B. Y. (2014). Hot-hand bias in rhesus monkeys. Journal of Experimental Psychology: Animal Learning and Cognition, 40(3), 280.

Blanchard, T. C., Hayden, B. Y., & Bromberg-Martin, E. S. (2015). Orbitofrontal cortex uses distinct codes for different choice attributes in decisions motivated by curiosity. Neuron, 85(3), 602–614.

Cai, X., & Padoa-Schioppa, C. (2014). Contributions of orbitofrontal and lateral prefrontal cortices to economic choice and the good-to-action transformation. Neuron, 81(5), 1140–1151.

Cavanagh, P., & Alvarez, G. A. (2005). Tracking multiple targets with multifocal attention. Trends in cognitive sciences, 9(7), 349–354.

Cisek, P. (2012). Making decisions through a distributed consensus. Current opinion in neurobiology, 22(6), 927–936.

Cohen, Y., Schneidman, E., & Paz, R. (2021). The geometry of neuronal representations during rule learning reveals complementary roles of cingulate cortex and putamen. Neuron, 109(5), 839–851.

Di Lollo, V. (2012). The feature-binding problem is an ill-posed problem. Trends in cognitive sciences, 16(6), 317–321.

Dong, Y., Mihalas, S., Qiu, F., von der Heydt, R., & Niebur, E. (2008). Synchrony and the binding problem in macaque visual cortex. Journal of vision, 8(7), 30–30.

Ebitz, R. B., & Hayden, B. Y. (2021). The population doctrine revolution in cognitive neurophysiology. arXiv preprint arXiv:2104.00145.

Elsayed, G. F., Lara, A. H., Kaufman, M. T., Churchland, M. M., & Cunningham, J. P. (2016). Reorganization between preparatory and movement population responses in motor cortex. Nature communications, 7(1), 1–15.

Farashahi, S., Azab, H., Hayden, B., & Soltani, A. (2018). On the flexibility of basic risk attitudes in monkeys. Journal of Neuroscience, 38(18), 4383–4398.

Farashahi, S., Donahue, C. H., Hayden, B. Y., Lee, D., & Soltani, A. (2019). Flexible combination of reward information across primates. Nature human behaviour, 3(11), 1215–1224.

Feierstein, C. E., Quirk, M. C., Uchida, N., Sosulski, D. L., & Mainen, Z. F. (2006). Representation of spatial goals in rat orbitofrontal cortex. Neuron, 51(4), 495–507.

Fine, J. M., & Hayden, B. Y. (2021). The whole prefrontal cortex is premotor cortex. arXiv preprint arXiv:2106.04651.

Fusi, S., Miller, E. K., & Rigotti, M. (2016). Why neurons mix: high dimensionality for higher cognition. Current opinion in neurobiology, 37, 66–74.

Gao, P., & Ganguli, S. (2015). On simplicity and complexity in the brave new world of large-scale neuroscience. Current opinion in neurobiology, 32, 148–155.

Ghose, G. M., & Maunsell, J. (1999). Specialized representations in visual cortex: a role for binding?. Neuron, 24(1), 79–85.

Hare, T. A., Schultz, W., Camerer, C. F., O’Doherty, J. P., & Rangel, A. (2011). Transformation of stimulus value signals into motor commands during simple choice. Proceedings of the National Academy of Sciences, 108(44), 18120–18125.

Hayden, B. Y., & Moreno-Bote, R. (2018). A neuronal theory of sequential economic choice. Brain and Neuroscience Advances, 2, 2398212818766675.

Hayden, B., Smith, D. V., & Platt, M. (2010). Cognitive control signals in posterior cingulate cortex. Frontiers in human neuroscience, 4, 223.

Hayden, B. Y. (2019). Why has evolution not selected for perfect self-control?. Philosophical Transactions of the Royal Society B, 374(1766), 20180139.

Hayden, B. Y., & Niv, Y. (2021). The case against economic values in the orbitofrontal cortex (or anywhere else in the brain). Behavioral Neuroscience, 135(2), 192.

Heilbronner, S. R., & Hayden, B. Y. (2016). The description-experience gap in risky choice in nonhuman primates. Psychonomic bulletin & review, 23(2), 593–600.

Heilbronner, S. R., & Hayden, B. Y. (2016). Dorsal anterior cingulate cortex: a bottom-up view. Annual review of neuroscience, 39, 149–170.

Holcombe, A. O., & Clifford, C. W. (2012). Failures to bind spatially coincident features: comment on Di Lollo. Trends in cognitive sciences, 16(8), 402.

Hunt, L. T., Dolan, R. J., & Behrens, T. E. (2014). Hierarchical competitions subserving multi-attribute choice. Nature neuroscience, 17(11), 1613–1622.

Hunt, L. T., & Hayden, B. Y. (2017). A distributed, hierarchical and recurrent framework for reward-based choice. Nature Reviews Neuroscience, 18(3), 172–182.

Hunt, L. T., Malalasekera, W. N., de Berker, A. O., Miranda, B., Farmer, S. F., Behrens, T. E., & Kennerley, S. W. (2018). Triple dissociation of attention and decision computations across prefrontal cortex. Nature neuroscience, 21(10), 1471–1481.

Kable, J. W., & Glimcher, P. W. (2007). The neural correlates of subjective value during intertemporal choice. Nature neuroscience, 10(12), 1625–1633.

Kable, J. W., & Glimcher, P. W. (2009). The neurobiology of decision: consensus and controversy. Neuron, 63(6), 733–745.

Kacelnik, A., Vasconcelos, M., Monteiro, T., & Aw, J. (2011). Darwin’s “tug-of-war” vs. starlings’“horse-racing”: how adaptations for sequential encounters drive simultaneous choice. Behavioral Ecology and Sociobiology, 65(3), 547–558.

Kawahara, J. I., & Yamada, Y. (2006). Two noncontiguous locations can be attended concurrently: Evidence from the attentional blink. Psychonomic bulletin & review, 13(4), 594–599.

Kennerley, S. W., Dahmubed, A. F., Lara, A. H., & Wallis, J. D. (2009). Neurons in the frontal lobe encode the value of multiple decision variables. Journal of cognitive neuroscience, 21(6), 1162–1178.

Kim, S., Hwang, J., & Lee, D. (2008). Prefrontal coding of temporally discounted values during intertemporal choice. Neuron, 59(1), 161–172.

Kimmel, D. L., Elsayed, G. F., Cunningham, J. P., & Newsome, W. T. (2020). Value and choice as separable and stable representations in orbitofrontal cortex. Nature communications, 11(1), 1–19.

Krajbich, I., Armel, C., & Rangel, A. (2010). Visual fixations and the computation and comparison of value in simple choice. Nature neuroscience, 13(10), 1292–1298.

Mante, V., Sussillo, D., Shenoy, K. V., & Newsome, W. T. (2013). Context-dependent computation by recurrent dynamics in prefrontal cortex. nature, 503(7474), 78–84.

Maisson, D. J., Yoo, S. B. M., Wang, M. Z., Cash-Padgett, T. V., Zimmermann, J., & Hayden, B. (2021). Differential encoding of safe and risky offers. bioRxiv.

McMains, S. A., & Somers, D. C. (2004). Multiple spotlights of attentional selection in human visual cortex. Neuron, 42(4), 677–686.

Niv, Y. (2019). Learning task-state representations. Nature neuroscience, 22(10), 1544–1553.

O’Neill, M., & Schultz, W. (2010). Coding of reward risk by orbitofrontal neurons is mostly distinct from coding of reward value. Neuron, 68(4), 789–800.

Padoa-Schioppa, C., & Assad, J. A. (2006). Neurons in the orbitofrontal cortex encode economic value. Nature, 441(7090), 223–226.

Parthasarathy, A., Tang, C., Herikstad, R., Cheong, L. F., Yen, S. C., & Libedinsky, C. (2019). Time-invariant working memory representations in the presence of code-morphing in the lateral prefrontal cortex. Nature communications, 10(1), 1–11.

Pastor-Bernier, A., Stasiak, A., & Schultz, W. (2019). Orbitofrontal signals for two-component choice options comply with indifference curves of Revealed Preference Theory. Nature communications, 10(1), 1–19.

Paxinos, G., Huang, X. F., & Toga, A. W. (2000). The rhesus monkey brain in stereotaxic coordinates.

Pearson, J., Hayden, B., & Platt, M. (2010). Explicit information reduces discounting behavior in monkeys. Frontiers in psychology, 1, 237.

Pezzulo, G., & Cisek, P. (2016). Navigating the affordance landscape: feedback control as a process model of behavior and cognition. Trends in cognitive sciences, 20(6), 414–424.

Platt, M. L., & Glimcher, P. W. (1999). Neural correlates of decision variables in parietal cortex. Nature, 400(6741), 233–238.

Raghuraman, A. P., & Padoa-Schioppa, C. (2014). Integration of multiple determinants in the neuronal computation of economic values. Journal of Neuroscience, 34(35), 11583–11603.

Rangel, A., Camerer, C., & Montague, P. R. (2008). A framework for studying the neurobiology of value-based decision making. Nature reviews neuroscience, 9(7), 545–556.

Raposo, D., Kaufman, M. T., & Churchland, A. K. (2014). A category-free neural population supports evolving demands during decision-making. Nature neuroscience, 17(12), 1784–1792.

Rich, E. L., & Wallis, J. D. (2016). Decoding subjective decisions from orbitofrontal cortex. Nature neuroscience, 19(7), 973–980.

Rigotti, M., Barak, O., Warden, M. R., Wang, X. J., Daw, N. D., Miller, E. K., & Fusi, S. (2013). The importance of mixed selectivity in complex cognitive tasks. Nature, 497(7451), 585–590.

Roesch, M. R., Taylor, A. R., & Schoenbaum, G. (2006). Encoding of time-discounted rewards in orbitofrontal cortex is independent of value representation. Neuron, 51(4), 509–520.

Roskies, A. L. (1999). The binding problem. Neuron, 24(1), 7–9.

Rustichini, A., & Padoa-Schioppa, C. (2015). A neuro-computational model of economic decisions. Journal of neurophysiology, 114(3), 1382–1398.

Samejima, K., Ueda, Y., Doya, K., & Kimura, M. (2005). Representation of action-specific reward values in the striatum. Science, 310(5752), 1337–1340.

Saxena, S., & Cunningham, J. P. (2019). Towards the neural population doctrine. Current opinion in neurobiology, 55, 103–111.

Shadlen, M. N., & Movshon, J. A. (1999). Synchrony unbound: a critical evaluation of the temporal binding hypothesis. Neuron, 24(1), 67–77.

Singer, W., & Gray, C. M. (1995). Visual feature integration and the temporal correlation hypothesis. Annual review of neuroscience, 18(1), 555–586.

Sleezer, B. J., Castagno, M. D., & Hayden, B. Y. (2016). Rule encoding in orbitofrontal cortex and striatum guides selection. Journal of Neuroscience, 36(44), 11223–11237.

Strait, C. E., Sleezer, B. J., Blanchard, T. C., Azab, H., Castagno, M. D., & Hayden, B. Y. (2016). Neuronal selectivity for spatial positions of offers and choices in five reward regions. Journal of neurophysiology, 115(3), 1098–1111.

Strait, C. E., Blanchard, T. C., & Hayden, B. Y. (2014). Reward value comparison via mutual inhibition in ventromedial prefrontal cortex. Neuron, 82(6), 1357–1366.

Strait, C. E., Sleezer, B. J., & Hayden, B. Y. (2015). Signatures of value comparison in ventral striatum neurons. PLoS Biol, 13(6), e1002173.

Tang, C., Herikstad, R., Parthasarathy, A., Libedinsky, C., & Yen, S. C. (2020). Minimally dependent activity subspaces for working memory and motor preparation in the lateral prefrontal cortex. Elife, 9, e58154.

Treisman, A. (1996). The binding problem. Current opinion in neurobiology, 6(2), 171–178.

Treisman, A. M., & Gelade, G. (1980). A feature-integration theory of attention. Cognitive psychology, 12(1), 97–136.

Tsujimoto, S., Genovesio, A., & Wise, S. P. (2009). Monkey orbitofrontal cortex encodes response choices near feedback time. Journal of Neuroscience, 29(8), 2569–2574.

Tucker, L. R. (1966). Some mathematical notes on three-mode factor analysis. Psychometrika, 31(3), 279–311.

Urai, A. E., Doiron, B., Leifer, A. M., & Churchland, A. K. (2021). Large-scale neural recordings call for new insights to link brain and behavior. arXiv preprint arXiv:2103.14662.

Von der Malsburg, C. (1995). Binding in models of perception and brain function. Current opinion in neurobiology, 5(4), 520–526.

Von der Malsburg, C. (1999). The what and why of binding: the modeler’s perspective. Neuron, 24(1), 95–104.

Wang, M. Z., Hayden, B., & Heilbronner, S. (2020). Anatomically distinct OFC-PCC circuits relay choice from value space to action space. bioRxiv.

Widge, A. S., Heilbronner, S. R., & Hayden, B. Y. (2019). Prefrontal cortex and cognitive control: new insights from human electrophysiology. F1000Research, 8.

Wolfe, J. M. (2012). The binding problem lives on: Comment on Di Lollo. Trends in cognitive sciences, 16(6), 307.

Wunderlich, K., Rangel, A., & O’Doherty, J. P. (2009). Neural computations underlying action-based decision making in the human brain. Proceedings of the National Academy of Sciences, 106(40), 17199–17204.

Yoo, S. B. M., Hayden, B. Y., & Pearson, J. M. (2021). Continuous decisions. Philosophical Transactions of the Royal Society B, 376(1819), 20190664.

Yoo, S. B. M., & Hayden, B. Y. (2020). The transition from evaluation to selection involves neural subspace reorganization in core reward regions. Neuron, 105(4), 712–724.

Yoo, S. B. M., & Hayden, B. Y. (2018). Economic choice as an untangling of options into actions. Neuron, 99(3), 434–447.

Yoo, S. B. M., Sleezer, B. J., & Hayden, B. Y. (2018). Robust encoding of spatial information in orbitofrontal cortex and striatum. Journal of cognitive neuroscience, 30(6), 898–913.

Zeki, S. (2020). “Multiplexing” cells of the visual cortex and the timing enigma of the binding problem. European Journal of Neuroscience.

Zou, H., & Hastie, T. (2003). Regression shrinkage and selection via the elastic net, with applications to microarrays. JR Stat Soc Ser B, 67, 301–20.

